# Autophagy suppresses microglial activation and enhances M2 polarization via the mTOR/ULK1 pathway after optic nerve crush

**DOI:** 10.64898/2026.06.11.731044

**Authors:** Hong-Ying Li, Xi Hong

**Author notes:** Corresponding Author: Hong-Ying Li, Medical Experimental Center, School of Basic Medical Sciences and Public Health, Jinan University, Guangzhou 510632, China. These authors contribute equally to the work.

## Abstract

**Purpose:** To investigate whether rapamycin can regulate microglial activation and polarization via mTOR and its downstream signals via autophagy both *in vivo* and *in vitro*.

**Methods:** The *in vivo* study used wild type C57BL/6 mice that were intraperitoneally injected with rapamycin (2 mg/kg) plus ONC. The BV2 cell line was used in the *in vitro* study and the cells were incubated with rapamycin (50 nM) or transfected with a specific mTOR-targeting small interfering RNA (si-mTOR). Immunohistochemical staining was used to observe the changes in the morphology and cell surface area of microglia and Weste blotting analysis was used for detection of the changes in the proteins related autophagy, microglia polarization and mTOR pathway after the retinal tissue or the cell samples were collected.

**Results:** These results indicate that rapamycin increases autophagy and M2 polarization by inhibiting p-mTOR in wild-type C57BL/6 mice *in vivo*. In the BV2 cell line, rapamycin and si-mTOR can enhance autophagy and promote M2 polarization by inhibiting the p-mTOR/p-Unc-51-like kinase 1 (p-ULK1) pathway.

**Conclusions:** In conclusion, this work contributes to the understanding of the complex interplay among rapamycin, autophagy and microglial activation/polarization, highlights the downstream signaling pathway of mTOR, and highlights the potential therapeutic effects of autophagy-modulating drugs in retinal neuroinflammation and neurodegeneration after TON.

## Introduction

The degeneration of retinal ganglion cells (RGCs) is the cause of vision decline in patients after traumatic optic neuropathy (TON). The mechanisms of retinal neurodegeneration under these conditions have been investigated for a long period of time by researchers worldwide. Among them, the role of neuroinflammation mediated by microglia has received much attention. Microglia are tissue macrophages and resident immune cells in the central nervous system (CNS) that originate from yolk sac primitive macrophages. Brain-resident microglia are involved in the maintenance of the neural microenvironment and inflammatory observation, playing a surveillance role in the CNS ^1,2^. Microglia can be activated by various pathological events, such as trauma ^3^, depression (Wang et al., 2022), neuroinflammation and neurodegeneration ^4^, and rapidly change their morphology and function (Nayak et al., 2014; Prinz et al., 2019; Vidal-Itriago et al., 2022). After activation, the microglial somas became larger, and the ramified processes became shorter and thicker ^5^. Additionally, activated microglia secrete various cytokines and signaling molecules and are polarized into two different phenotypes: M1 and M2 ^6^. M1 is the classical proinflammatory phenotype, which causes tissue inflammation or cell death and the secretion of inflammatory factors such as interleukin-1β (IL-1β), IL-6, inducible nitric oxide synthase (iNOS), and tumor necrosis factor alpha (TNF-α); the M2 phenotype (also known as the alternative phenotype), which is recognized as an anti-inflammatory phenotype and participates in inflammatory suppression and tissue remodeling, expresses anti-inflammatory factors, including arginase-1 (Arg-1), cluster of differentiation 206 (CD206), and IL-10 ^7–11^. These two different phenotypes are interconvertible in different stages of disease, making it possible for microglia to accommodate specific circumstances ^12^. Although some researchers believe that it is somewhat outdated to categorize microglia as ‘‘resting versus activated’’ or ‘‘M1 versus M2’’ ^6^; however, these kinds of classifications can still help understand the phenotype and function of microglia under various circumstances, such as brain trauma or neurodegenerative diseases, and are chosen by multiple biologists to date ^13–16^. Therefore, this type of nomenclature was still applied in this study.

Autophagy is an indispensable metabolic pathway in eukaryotic development and growth and is involved in a multitude of physiological and pathological processes ^17,18^. Autophagy can be activated to degrade organelles, proteins and metabolites to recycle and reuse cell components ^19^. Growing evidence suggests that autophagy is involved in microglial activation and polarization. mTOR is a serine/threonine kinase that plays a crucial role in cell growth and metabolism. There are two distinct mTOR complex forms: mTOR complex 1 (mTORC1) and mTOR complex 2 (mTORC2) ^20^. mTORC1 is a multiprotein complex that plays a central role in regulating cell growth, metabolism, and autophagy in response to various stimuli. The composition of mTORC1 includes the following key components: mTOR (a serine/threonine kinase), regulatory-associated protein of mTOR (RAPTOR) and mammalian lethal with SEC13 protein 8 (mLST8). mTORC1 is a key regulator of autophagy, and once activated, mTORC1 further promotes anabolic processes while inhibiting autophagy through the modulation of ULK1 ^21^. MTORC1 is sensitive to rapamycin, which is an mTOR inhibitor, and the mechanism by which rapamycin activates autophagy involves the formation of a complex with FK506-binding protein 12 (FKBP12) and binding to the FKBP12-rapamycin binding domain of mTOR (FRB), leading to the inhibition of mTORC1 activity; on the other hand, mTOR complex 2 (mTORC2) is not sensitive to acute treatment with rapamycin ^22,23^.

Some studies have shown that rapamycin or mTOR can change the state of microglial activation/polarization; however, most of these studies focused only on a limited scope of this field. For example, they performed only *in vivo* or *in vitro* studies but not both; alternatively, they focused on the regulation of microglial activation but not polarization at the same time ^24–28^. Therefore, it is important to design a thorough study that includes both *in vivo* and *in vitro* experiments to examine the roles of rapamycin and mTOR at the same time and to investigate the activation and polarization of microglia synchronously to understand the complex relationships among rapamycin, autophagy, mTOR and microglia. In this study, we used C57BJ/6 mice treated with ONC and the BV2 cell line to study the underlying mechanism more clearly.

## Materials and methods

### Animals

C57BL/6 male mice were purchased from Guangdong Medical Laboratory Animal Center (Guangdong, China). These animals were maintained under specific-pathogen-free (SPF) conditions on a 12-hour light/dark cycle. The room temperature was automatically maintained at 21–25°C with a relative humidity of 45–65%. Food and water were provided ad libitum. All these animal studies were performed in accordance with the guidelines of the Animal Ethics Committee of Jinan University, Guangzhou, Guangdong Province, China.

C57BL/6 male mice (18–20 g) aged 6–8 weeks were randomly assigned to the following five groups: the normal group (n = 7), the group with optic nerve crush (abbreviated as the ONC group: n = 8), the ONC + rapamycin (2 mg/kg) group (n = 7), and the ONC + 0.4% dimethyl sulfoxide (DMSO) group (n = 8). In each group, 3–4 mice were used for the preparation of whole-mounted proteins, and the remainder of each group was prepared for protein analysis via Western blotting analysis after euthanasia.

### Preparation and usage of rapamycin

The powder of rapamycin (powder, MCE, HY10219) was dissolved in DMSO to a concentration of 20 mg/mL as the stock solution. Before the daily injection, the stock solution was diluted to a final concentration of 0.08 mg/mL with sterilized phosphate-buffered saline (PBS, pH 7.4) and mixed well. Each mouse weighed 18–20 g, and the dosage was 2 mg/kg (rapamycin) for each mouse. A volume of 0.5 mL was injected into each mouse daily. In the control group, an equal volume of 0.4% DMSO in PBS was injected. Before each injection, the solution was mixed well, and the liquid was injected into the abdominal cavity without damaging the organs in the abdominal cavity with proper operation. The intraperitoneal injection of rapamycin or vehicle was accomplished with a sterilized 1 ml syringe. The injection lasted from two days before ONC to 2 days after ONC; therefore, a total of five injections were completed (Fig. 1A).

**Fig. 1.**
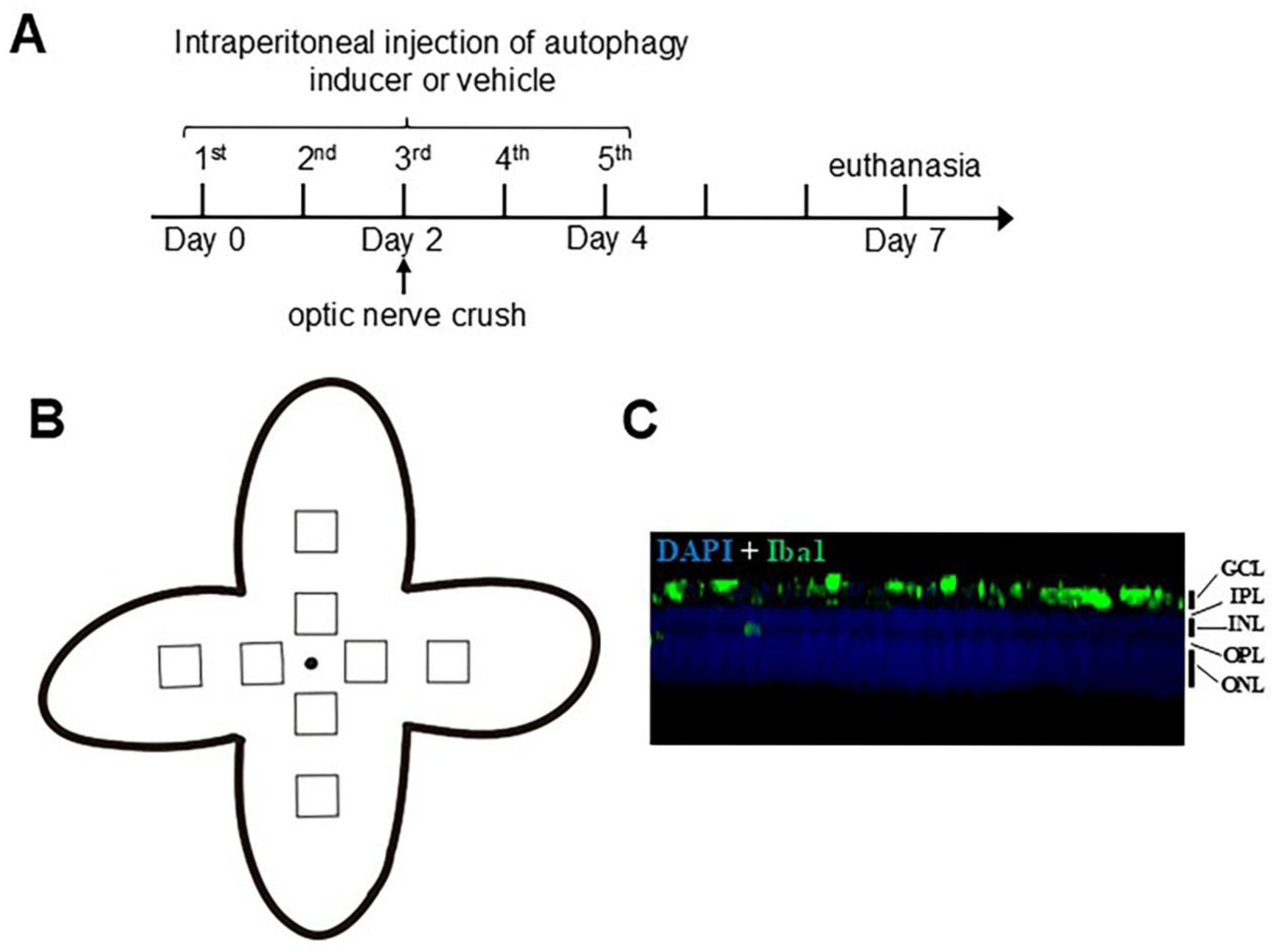
The figures related to the method part. **A** The *in vivo* experimental design. Experimental timeline: An autophagy inducer or vehicle was administered via intraperitoneal injection on days 0, 2, 4, and 6. Optic nerve crush was performed on day 2, and euthanasia was carried out on day 7. **B** The retinal whole-mount sampling method. Schematic representation of a mouse retinal flat mount: the retina was sectioned into a cloverleaf shape, with two square regions selected for imaging in each quadrant. The first image was taken from the edge of the optic disc, and the second image was captured 800 μm away from the optic disc. Each image had dimensions of 400 μm × 400 μm. **C** The distribution of Iba1-positive microglia in the retina along the Z axis. Confocal fluorescence microscopy of retinal layers: DAPI staining (blue) was used to label cell nuclei, while Iba1 staining (green) highlighted microglia. The retinal layers were distinguishable, with microglia distributed in three layers: the ganglion cell layer (GCL), inner plexiform layer (IPL), and outer plexiform layer (OPL). Only microglia in the GCL were selected for confocal imaging.

### The preparation of Avertin

To make 1.25% Avertin solution, 1.25 g powder of tribromethyl alcohol (TBE, Sigma-Aldrich, T48402) was dissolved in 100 ml solution in a light shield bottle by adding 2.5 ml tertiary amyl alcohol (Sigma Aldrich, CAS: 75-85-4) and double distilled water in sequence. The solution was then blended by magnetic blender overnight till TBE was fully dissolved. Then the solution was filtered through a 0.22 μm filter and stored in 4°C for up to 1 week.

### Optic nerve crush surgery

The surgery was performed via a similar procedure as the published procedure adopted by Tang et al. ^29^. In brief, the mice were deeply anesthetized (Avertin, 20 μl/g, i.p.), eye drops of oxybuprocaine (Santen Pharmaceutical Co., Ltd.) were used to anesthetize the cornea locally. An incision was made at the conjunctiva of the left eye at approximately the four o’clock position via a pair of spring scissors, and the optic nerve was located after dissecting the connective tissue and the extraocular muscles. The optic nerve (ON) was crushed for 10 seconds at 0.5 mm from the optic disk. Tobramycin eye ointment was used on the eye surface after surgery to prevent infection. After surgery, the eyes of the mice were observed every day until euthanasia to ensure that the vessels had not been damaged.

### Immunofluorescence staining

After the mice were euthanized at the set timepoint, the eyes were removed from the orbit and then immersed in 4% paraformaldehyde (PFA, Sigma, CAS 30525-89-4) for 1 hour to ensure fixation. After the cornea, lens and vitreous body were removed, the retinas were collected and rinsed thoroughly with chilled 0.01 M PBS (pH 7.4). The retinas were subsequently divided into four cuts into a four-leaf clover configuration (Fig. 1B). After nonspecific binding sites were blocked by incubating the retinas with 10% bovine serum albumin (BSA) for 1 hour, the samples were incubated with primary antibody against ionized calcium binding adaptor molecule 1 (Iba1, WAKO, 1:1000) for 36 hours to target specific antigens. After being washed with 0.01 M PBS (pH 7.4), the retinas were then incubated with an Alexa Fluor^TM^ 488-conjugated goat anti-rabbit secondary antibody (Invitrogen, 1:1000) for 2 hours at room temperature to visualize the primary antibody binding. After that, the retinas were washed with 0.01 M PBS (pH 7.4) six times (10 min each time), and following this step, the retinas were incubated with 4’,6-diamidino-2-phenylindole (DAPI, 1:2000, Electron Microcopy Sciences, USA) for 30 min at room temperature for nuclear counterstaining. Finally, the retinas were mounted with Fluor-Mount medium (Sigma, F4680-25) for microscopic analysis after they were rinsed with 0.01 M PBS (pH 7.4) six times.

### Retinal microglial cell capture

The capture of microglia stained with Iba1 (green) in the whole ganglion cell layer (GCL) of the mouse retina was achieved via Z-stack scanning via a confocal microscope (Carl Zeiss LSM700, Oberkochen, Germany). The GCL was distinguished from the other layers by the specific arrangement of one-layer nuclei, which were stained with DAPI (Fig. 1C). For each leaf of the four-leaf-clover-shaped retina, two images with a size of 400 μm × 400 μm were captured either adjacent to the edge of the optic disc or 0.8 mm away from the first image at 200× magnification; finally, eight images per retina were captured (Fig. 1B). The scanning interval for Z-stacking is 0.69 μm. ImageJ software (version 1.41) was used to merge all the images acquired by scanning at one location into a completely stacked final picture, and the area occupied by microglial somas in the image was measured. The average Iba1-positive area (μm^2^) of the microglial cell bodies per mm^2^ was obtained from the 8 regions in each animal. The average positive area (μm^2^) obtained from all animals in each group was used for comparison between groups.

### Retinal protein extraction

For the animal experiments, after the mice were euthanized by breaking the neck, the retinas were isolated in chilled 0.01 M PBS (pH 7.4) and put into radioimmunoprecipitation assay (RIPA) lysis buffer (Thermo Scientific, 89900) supplemented with phosphatase inhibitor (PPI, Thermo Scientific, 78440) and protease inhibitor (PI, Thermo Scientific, 78440). An ultrasonic cell disruptor (Sonics & Materials) was used to break the cells, and the suspension was centrifuged at 11000 revolutions per minute (RPM) for 10 minutes after 30 minutes of quiescence in a centrifuge tube on ice. The supernatant was collected and stored at -80°C until use for Western blot analysis.

### BV2 cell culture and rapamycin treatment

The BV2 cell line was a gift from Professor Kwok-Fai So in Jinan University, Guangzhou, China. The BV2 cell line represents an immortalized mouse microglial cell line that retains a substantial portion of the morphological, phenotypic, and functional attributes of primary microglia. BV2 cells were seeded into T50 flasks and maintained in culture medium consisting of 10% fetal bovine serum (FBS; Gibco 10099141)-Dulbecco’s modified Eagle’s medium (DMEM; Gibco 11965092) under humid conditions with 5% CO_2_ and 95% air at 37°C. The culture medium was replaced with fresh medium every two days to ensure optimal growth conditions. Once the cells reached approximately 80% confluence in the T50 flasks, they were enumerated and transferred to a 6-well cell plate until they adhered to the wall approximately 12 hours later. For the rapamycin (MCE, HY-10219) treatment, the stock solution of 20 mg/mL rapamycin was diluted to a concentration of 50 nM in each well with 2 × 10^5^ BV2 cells as well as 2 ml of 10% FBS-DMEM. The cells were incubated with rapamycin for 6 hours. Moreover, the control group was incubated with 10% FBS-DMEM for an equivalent period.

### Small interfering RNA (siRNA)-mediated mTOR knockdown in BV2 cells

In a 6-well culture plate, 2 × 10^5^ BV2 cells were seeded into each well, which contained 2 mL of 10% FBS-DMEM. The cells were then incubated overnight in a humidified incubator maintained at 37°C with 5% CO_2_ and 95% air. Lipofectamine 2000 (Invitrogen, 19778030), si-mTOR (Santa Cruz, sc-35410), and non-silencing siRNA (Santa Cruz, sc-37007) were diluted separately in Opti-MEM (Gibco, 31985062). The diluted Lipofectamine 2000 was subsequently combined with both the diluted si-mTOR and the diluted non-silencing siRNA, resulting in final concentrations of 100 nM for both the si-mTOR mixture and the non-silencing siRNA mixture. The cells were then treated with the respective siRNA mixtures for 6 hours. After incubation, the siRNA mixtures were aspirated, and fresh 10% FBS-DMEM was added to the cells for an additional 48-hour incubation period. Each experimental group consisted of four wells to ensure reproducibility.

### BV-2 cell protein preparation

For the cellular experiment, following aspiration of the culture medium and rinsing with chilled 0.01 M PBS (pH 7.4), RIPA lysis buffer containing PI and PPI was introduced into the wells to facilitate cell lysis. The cells were subsequently harvested via a cell scraper and disrupted via an ultrasonic cell disruptor. Following a quiescence period of 30 minutes, the suspension was centrifuged at 11000 RPM for 10 minutes. The resulting supernatant was then collected and stored at -20°C for subsequent utilization.

### Western blotting analysis

Retinal protein and BV2 cell protein samples were quantified with a Pierce BCA protein assay kit (Beyotime, P0012s) and then incubated with sample loading buffer (Beyotime, P0015) at 95°C for 10 minutes. For electrophoresis, 12 micrograms of total protein from each sample were loaded per lane and separated initially at 80 V for 30 minutes, followed by 110V for 100 minutes. The samples were subsequently transferred onto polyvinylidene fluoride (PVDF) membranes (Bio-Rad, 1704157) via electrophoretic transfer at 300 milliamperes for 110 minutes. The membranes were then blocked with 5% BSA (Sigma, V900933) for 1 hour at room temperature. After blocking, the membranes were incubated overnight at 4°C with various primary antibodies specific to the target proteins, as outlined in Table 1. The membranes were washed with Tris-buffered saline containing Tween-20 (TBST, pH 7.5) after antibody incubation. Following the removal of the primary antibodies, the membranes were washed again and incubated with horseradish peroxidase (HRP) -conjugated secondary antibodies for 2 hours at room temperature. After washing, the membranes were treated with an HRP substrate (Millipore, WBLUF0500) to facilitate protein detection. Visualization of the membranes was achieved via the UVItec imaging system (Alliance 4.7), and quantitation was performed via the ImageJ program (version 1.41). Detailed information regarding the primary antibodies used in this study is provided in Table 1.

**Table 1.** Primary antibody information.

| Antibody | Brand | Product<br>Number | Working<br>Concentration |
| --- | --- | --- | --- |
| mTOR | Abcam | ab134903 | 1:4000 |
| p-mTOR | Abcam | S2248 | 1:2000 |
| ULK1 | CST | 14202 | 1:1000 |

| **Antibody** | **Brand** | **Product**<br>**Number** | **Working**<br>**Concentration** |
| --- | --- | --- | --- |
| p-ULK1 (Ser757) | CST | 8054 | 1:1000 |
| Beclin-1 | CST | 3495 | 1:1000 |
| p62 | CST | 23214 | 1:1000 |
| LC3A/B | CST | 12741 | 1:1000 |
| iNOS | Abcam | ab15323 | 1:250 |
| IL-6 | CST | 12912 | 1:1000 |
| Arg-1 | CST | 93668 | 1:1000 |
| CD206 | Abcam | ab64693 | 1:500 |
| Iba1 | WAKO | 20001 | 1:1000 |
| β-actin | CST | 9819 | 1:8000 |
| GAPDH | CST | 5174 | 1:8000 |
| Rabbit HRP-conjugated<br>secondary antibody | Jackson<br>Immuno<br>Research | 323-005-021 | 1:5000 |
| Mouse HRP-conjugated<br>secondary antibody | Jackson<br>Immuno<br>Research | 200-002-211 | 1:5000 |

### Statistics

All the data were analyzed via an unpaired t test, and the error bars represent the means ± standard errors (SEs). Statistical graphs were generated and plotted with GraphPad Prism 8.2.1 (GraphPad Software, Inc.). Statistical results are expressed as * *p* <0.05, ** *p* < 0.01, *** *p* < 0.001, **** *p* < 0.0001, NS = not significant. The data were considered different when *p* < 0.05.

## Results

### Effects of ONC on retinal autophagy and microglial cell activation/polarization

Five days after ONC, the retinal expression levels of LC3-II, p62 and Beclin-1 were similar to those in normal mouse retinas (*p* > 0.05) (Fig. 2A, B). Additionally, the Iba1-positive microglia changed from a resting phenotype, which had a ramified morphology with a small body and several long, highly branched processes, to an activated phenotype, which had a more rounded appearance with larger bodies and retracted and shorter processes (Fig. 2C, D). ImageJ software was used to quantify the positive area (µm^2^) of the microglial soma per mm^2^ in the retina, and the results showed that the average positive area of the microglial soma per mm^2^ in the normal retina was 5778 ± 758 µm^2^ and the average positive area of the microglial soma per mm^2^ in the retina 5 days after ONC was 40840 ± 1397 µm^2^; therefore it revealed an intensive increase in the mouse retina after ONC compared with the normal retina (Fig. 2C, D) (*p* < 0.0001). Moreover, western blot analysis revealed that the expression levels of the M1 microglial cell-related marker IL-6 (*p* < 0.05) and the M2 microglial cell-related marker Arg-1 (*p* < 0.01) increased in the ONC mouse retina (Fig. 2E, F), indicating that the numbers of both M1 and M2 microglia increased.

**Fig. 2.**
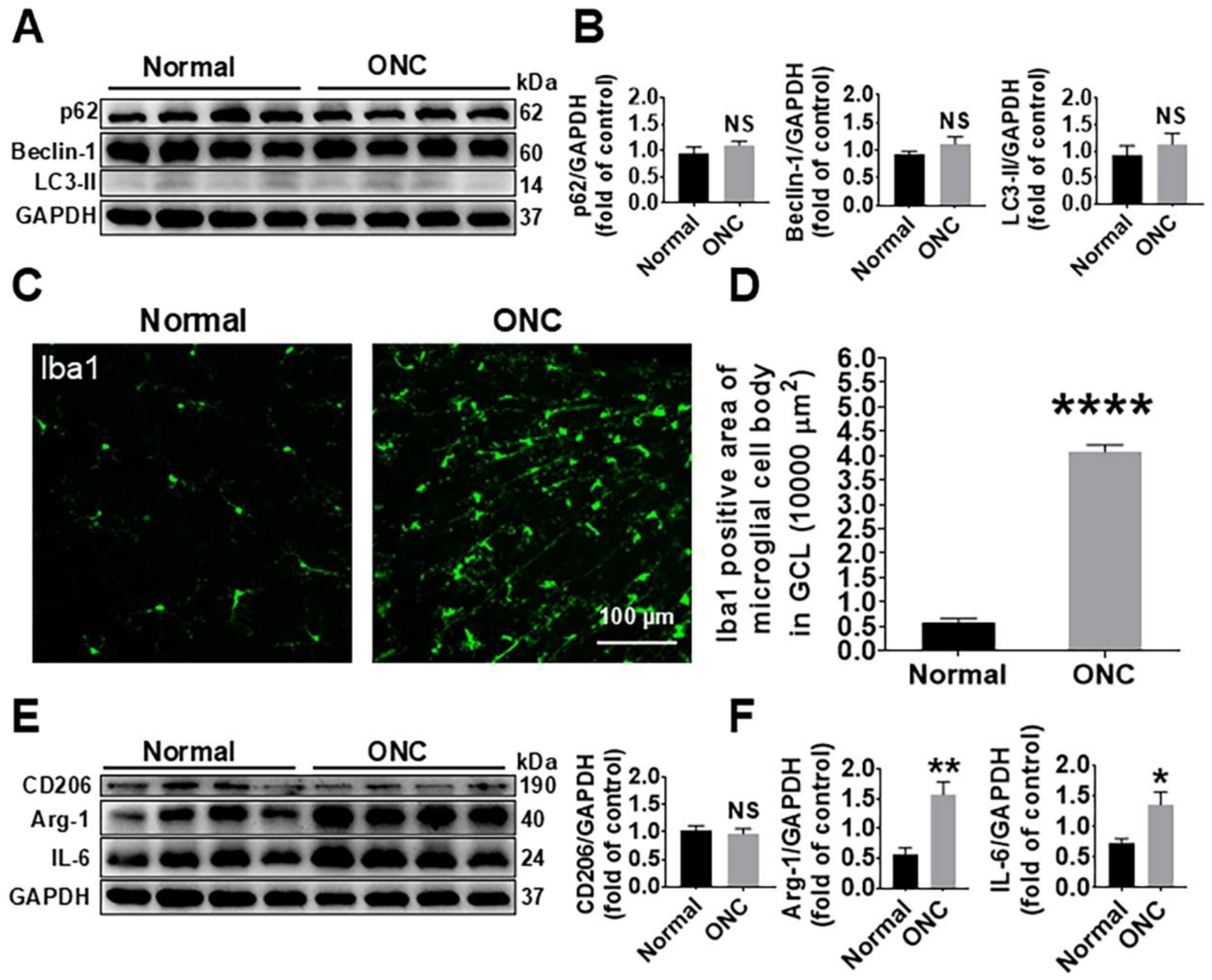
Effects of optic nerve crush (ONC) on autophagy and microglia activation in the retina. **A** Western blot analysis of p62, Beclin-1, LC3-II, and GAPDH in the retinas of the normal and ONC groups. N = 4. **B** p62, Beclin-1, and LC3-II expression levels were normalized to those of GAPDH in the normal and ONC groups. The data are presented as the fold change relative to the first band of the normal group. N = 4. **C** Immunofluorescence staining of Iba1 in the ganglion cell layer (GCL) of the normal and ONC groups. Scale bar = 100 μm. **D** Quantitative analysis of Iba1-positive microglial cell body area in the GCLs of the normal and ONC groups. The data are presented as the fold change relative to the first band of the normal group. N = 3 in the normal group and N = 4 in the ONC group. **E** Western blot analysis of CD206, Arginase-1, IL-6, and GAPDH in the normal and ONC groups. N = 4. **F** Quantitative analysis of CD206, Arginase-1, and IL-6 expression levels normalized to those of GAPDH in the normal and ONC groups. N = 4. The data are presented as the fold change relative to the first band of the normal group. * *p* < 0.05, ** *p* < 0.01, NS, not significant. The data are presented as the means ± SEMs.

### Rapamycin decreased microglial cell activation and drove the M2 polarization of microglia in the mouse retina after ONC

The western blot results revealed that rapamycin decreased the p-mTOR/mTOR ratio (*p* < 0.01) in the mouse retina, indicating that the intraperitoneal injection of 2 mg/kg rapamycin was suitable for activating autophagy in mice (Fig. 3A, B). The expression levels of LC3-II (*p* < 0.05) and beclin-1 (*p* < 0.01) were increased, whereas that of p62 was decreased (*p* < 0.001) (Fig. 3A, B). The Iba1-positive area of the microglial soma in the mouse ganglion cell layer (GCL) was 35250 ± 2026 µm^2^ per mm^2^ five days after ONC without rapamycin treatment and was 23780 ± 2486 µm^2^ per mm^2^ five days after ONC with rapamycin treatment, which meant that rapamycin treatment inhibited the activation of microglia (*p* < 0.05) (Fig. 3C, D). Moreover, the expression levels of M2 microglial cell markers, such as CD206 (*p* < 0.05) and Arginase-1 (*p* < 0.05), were increased, whereas the expression level of the M1 microglial marker IL-6 (*p* > 0.05) was not changed in the rapamycin-treated mouse retina (Fig. 3E, F), indicating that autophagy activation induced by rapamycin led to the microglial phenotype conversion to M2.

**Fig. 3.**
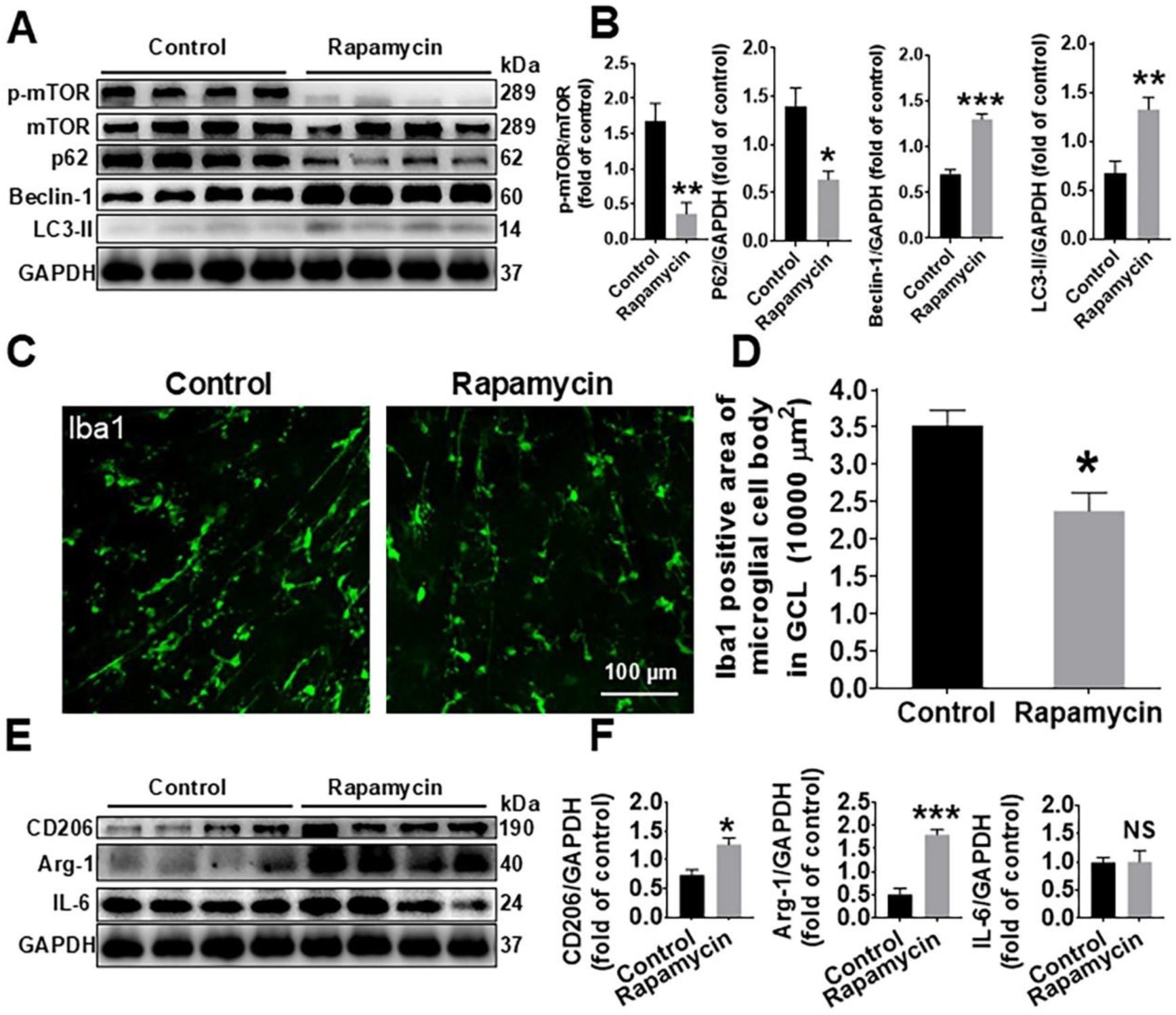
Effects of rapamycin on autophagy and microglial activation and polarization in the mouse retina. **A** Western blot analysis of autophagy-related proteins (p-mTOR/mTOR, p62, Beclin-1, and LC3-II) in the mouse retina, with GAPDH as a loading control. N = 4. **B** Quantitative analysis of the protein expression levels shown in panel A. **C** Immunofluorescence staining of Iba1-positive microglia in the granular cell layer (GCL) of the retina. Scale bar = 100 μm. N = 4 in the control group and N = 3 in the rapamycin group. **D** Quantitative analysis of the microglial cell body area shown in panel C. N = 4. **E** Western blot analysis of M1 and M2 microglial polarization markers (CD206, Arg-1, and IL-6) in the mouse retina, with GAPDH as a loading control. N = 4. **F** Quantitative analysis of the protein expression levels shown in panel E. N = 4. The data are presented as the means ± SEMs. \**p* < 0.05, \*\**p* < 0.01, \*\*\**p* < 0.001 compared with the control group. NS, not significant.

### BV-2 cells were polarized to the M2 phenotype after rapamycin treatment

BV-2 cells were incubated with rapamycin for 6 hours. According to the western blot results, the ratios of p-mTOR/mTOR (*p* < 0.05) and p-ULK1 (Ser757)/ULK1 (*p* < 0.01) were decreased in the rapamycin-treated BV-2 cells (Fig. 4A, B). In addition, the expression levels of LC3-II (*p* < 0.05) and beclin-1 (*p* < 0.01) were increased (Fig. 4A, B). Further experiments revealed that M2 microglial markers, such as CD206 (*p* < 0.01) and Arginase-1 (*p* < 0.05), were increased, whereas the M1 microglial marker iNOS was not changed (*p* > 0.05) (Fig. 4C, D).

**Fig. 4.**
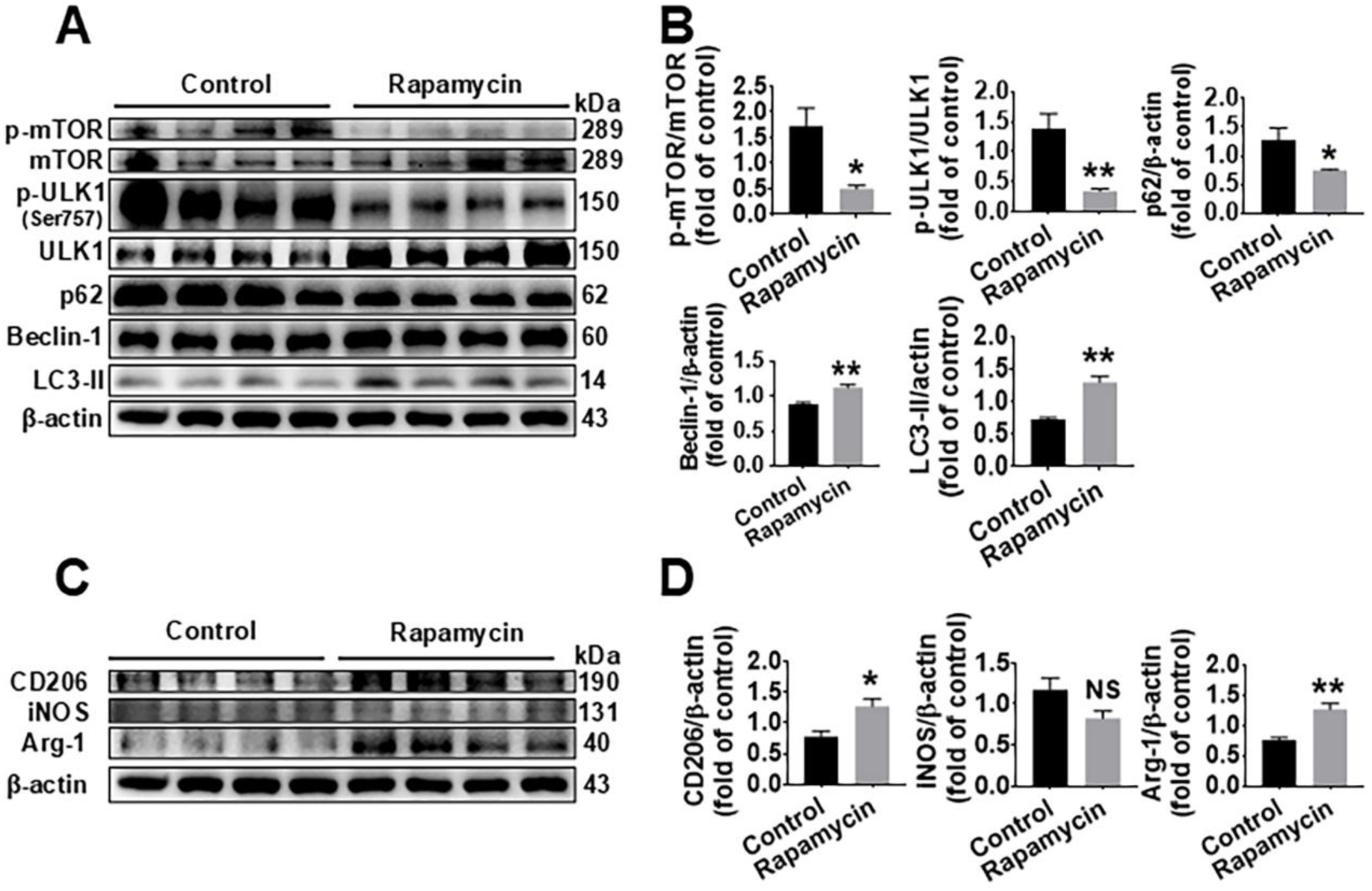
Effects of rapamycin on the expression of autophagy-related and M1/M2 microglial polarization proteins *in vitro*. **A** Western blot analysis of autophagy-related proteins (p-mTOR, mTOR, p-ULK1 (Ser757), ULK1, p62, Beclin-1, and LC3-II) in control and rapamycin-treated BV-2 cells, with β-actin as a loading control. N = 4. **B** Quantitative analysis of the Western blot results shown in panel A. The ratios of p-mTOR/mTOR, p-ULK1/ULK1, p62, Beclin-1, and LC3-II were normalized to those of β-actin and are expressed as the fold change relative to the control group. N = 4. **C** Western blot analysis of M1/M2 microglial polarization markers (CD206, iNOS, and Arg-1) in control and rapamycin-treated cells, with β-actin as a loading control. N = 4. **D** Quantitative analysis of the Western blot results shown in panel C. The expression levels of CD206, iNOS, and Arginase-1 (Arg-1) were normalized to those of β-actin and are expressed as the fold change relative to those in the control group. Each group consisted of four replicates. The “fold control” refers to the expression level of the first well in the control group. N = 4. Statistical analysis was performed via one-way ANOVA followed by Tukey’s post hoc test. The data are presented as the means ± SEMs. *p < 0.05, **p < 0.01 compared with the control group. NS, not significant.

### mTOR siRNA transfection promoted the polarization of BV-2 cells to the M2 phenotype

According to the western blot results, the ratios of both p-mTOR/mTOR (*p* < 0.01) and p-ULK1 (Ser757)/ULK1 (*p* < 0.001) were decreased in the mTOR siRNA-transfected BV-2 cells (Fig. 5). The expression levels of both LC3-II (*p* < 0.01) and beclin-1 (*p* < 0.05) were increased (Fig. 5). Moreover, the expression level of the M2 microglial marker CD206 was increased (*p* < 0.01) in the mTOR siRNA-transfected BV-2 cells (Fig. 5).

**Fig. 5.**
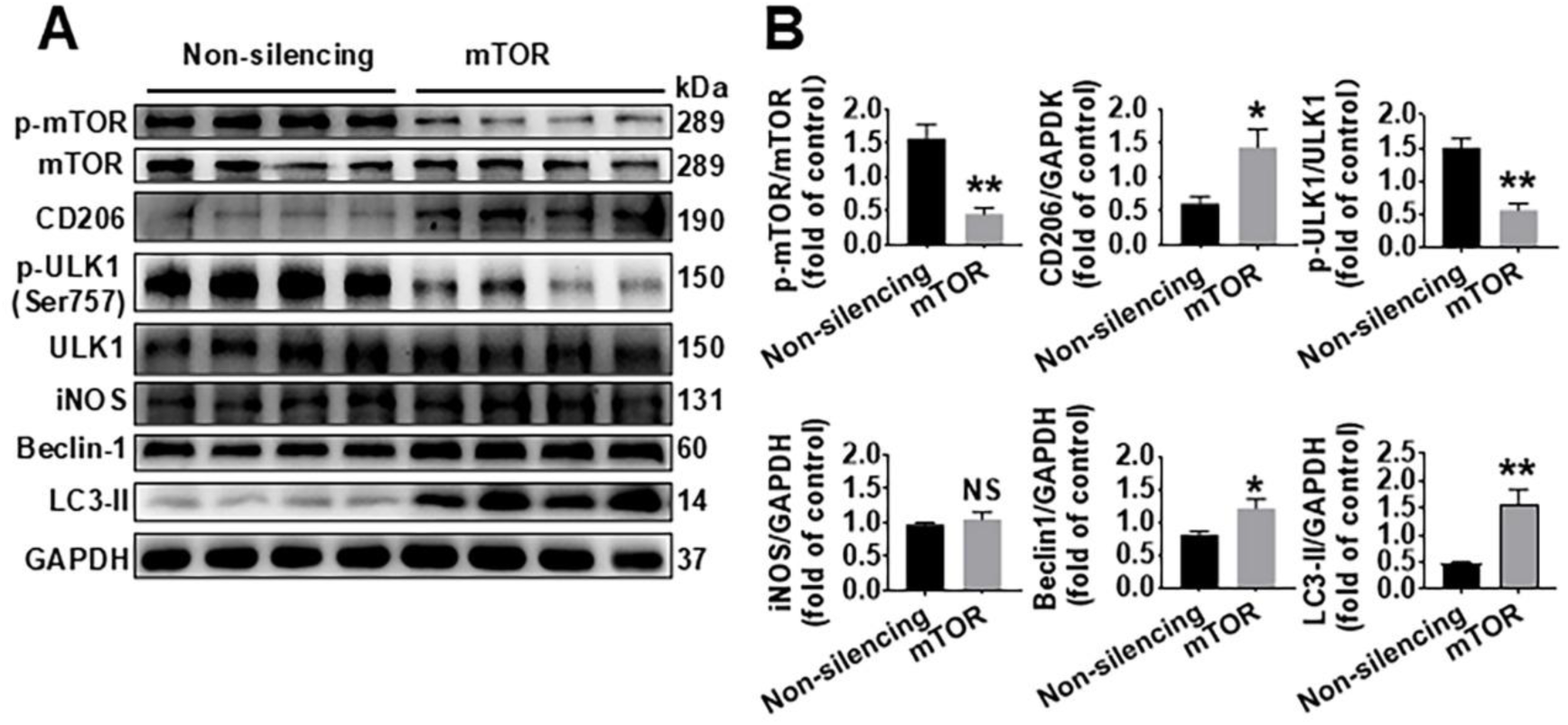
Effects of siRNA-mediated mTOR silencing on autophagy-related and microglial phenotype-related protein expression levels *in vitro*. **A** Western blot analysis of protein expression levels under nonsilencing and mTOR-silenced conditions via siRNA. The proteins analyzed included p-mTOR, mTOR, CD206, p-ULK1 (Ser757), ULK1, iNOS, Beclin-1, and LC3-II, with GAPDH as a loading control. The molecular weights (kDa) of the proteins are indicated on the right. N = 4. **B** Quantitative analysis of protein expression levels relative to those of the control (non-silencing condition). The graphs show the fold changes in the protein expression levels of p-mTOR/mTOR, CD206, p-ULK1/ULK1, iNOS, Beclin-1, and LC3-II. The data are presented as the means ± SEMs. N = 4. ∗*p* <0.05, ∗∗*p* <0.01. mTOR: mammalian target of rapamycin; p-ULK1 (Ser757): phosphorylated Unc-51-like kinase 1 at Serine 757; ULK1: Unc-51-like kinase 1; iNOS: inducible nitric oxide synthase; GAPDH: glyceraldehyde 3-phosphate dehydrogenase

## Discussion

Previous studies have demonstrated that RGC degeneration after TON might be correlated with microglial activation ^30–32^. In other words, the study of microglial activation and polarization may be a potential direction for investigating methods or drugs for treating TON. Microglia can be polarized into two different phenotypes: M1 proinflammatory and M2 anti-inflammatory microglia ^16,33^. Although there are some doubts about this dichotomous classification of microglial polarization ^6^, this categorization can still provide researchers with a simple and easy-to-understand way to distinguish different types of microglia with distinct characteristics and functions under complex circumstances and to find promising treatments for different diseases involving the activation of microglia/macrophages ^8,34,35^. More experiments, such as integrative analyses using single-cell technologies and multiomics ^36–38^, can be performed in the future to further classify specific cell states.

Autophagy has been shown to increase the M2 polarization of microglia in different situations, such as cerebral ischemia/reperfusion ^39,40^, neuroinflammation ^41^ and sepsis-associated encephalopathy ^42^. Rapamycin is a drug that can inhibit mTOR to increase autophagy *in vivo* and *in vitro*; therefore, it is widely used in studies related to autophagy. In this study, rapamycin was also used to increase autophagy in C57BL/6J mice and in the BV2 cell line. Our results showed that rapamycin could increase the expression levels of LC3 II and beclin-1 and downregulate the expression levels of p62 both *in vivo* and *in vitro*, which indicated that autophagy was activated after rapamycin treatment. Therefore, the dosage of rapamycin we used in this study was suitable for inducing autophagy both *in vivo* and *in vitro*. Furthermore, rapamycin was shown to increase the M2 marker CD206 and decrease the M1 marker iNOS, which indicated that the autophagy induced by rapamycin drove polarization to the M2 type.

How does autophagy influence microglial activation and polarization? According to a recent study, lipopolysaccharide (LPS) inhibits autophagy in BV-2 cells, which may lead to an inflammatory response and microglial polarization to the M1 type via the phosphorylation of ULK1 at Ser757 ^43^. ULK1 is a critical protein of the initial autophagosome formation complex, and its phosphorylation at Ser757 may suppress autophagy ^44–47^. The phosphorylation of ULK1 (Ser757) was inhibited in the *in vitro* experiments in this study, both in the mTOR siRNA-transfected BV-2 cells and in the rapamycin-treated BV-2 cells, indicating that rapamycin induces M2 polarization by regulating the mTOR/ULK1 pathway.

In conclusion, our study reveals the modulatory effects of autophagy induced by rapamycin and the downstream mTOR-ULK1 pathway in the retina and in BV2 cells (Fig. 6). Our study may provide new insights into the treatment of neurodegeneration after TON by influencing microglial activation and polarizatio

**Fig. 6.**
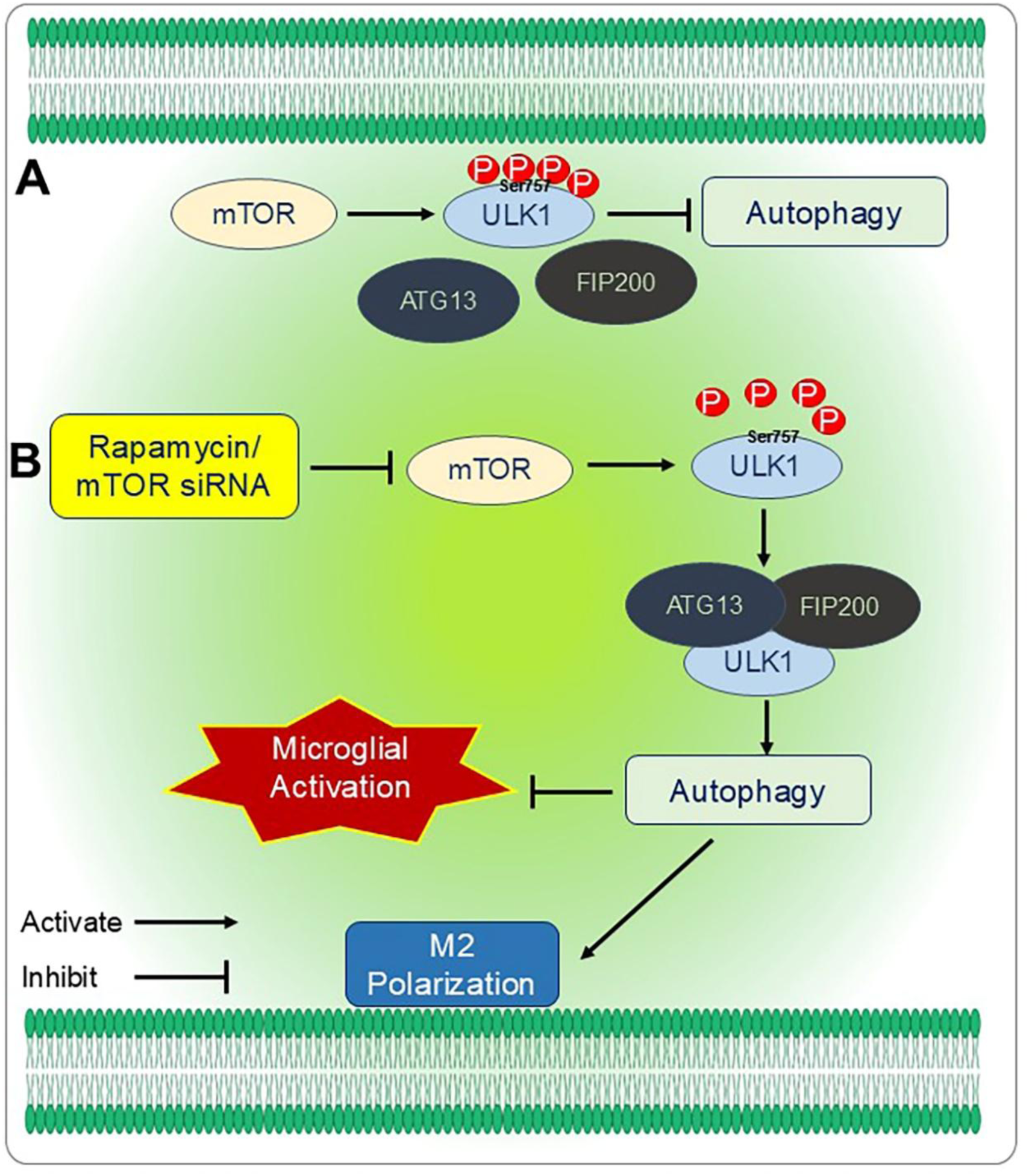
The potential mechanism by which autophagy affects microglial activation and polarization.

## Acknowledgments

I would like to express my gratitude to Guangdong-Hongkong-Macau Institute of CNS regeneration and Medical Experimental Center, School of Basic Medical Sciences and Public Health, Jinan University, Guangzhou, China for providing laboratory space and equipment to complete the experiments.

## Funding

This project was supported by the National Natural Science Foundation of China, under grant No. 81501091, awarded to Hong-Ying Li and the Natural Science Foundation of Guangdong Province, under grant No. 2015A030310201, awarded to Hong-Ying Li.

## Author Contributions

Hong-Ying Li: conceived the study, coordinated the project, designed the experiments, analyzed the data and revised the manuscript; Xi Hong: performed the experiments and prepared the first draft of the manuscript. Both authors read and approved the final manuscript. Hong-Ying Li and Xi Hong contributed equally to this work.

## Ethics Approval and Consent to Participate

All experimental procedures described in this article were reviewed and approved by the Ethics Committee of Jinan University.

## Availability of Data and Materials

All datasets used and/or analyzed during this study are available from the corresponding author on reasonable request.

## Disclosure

**Hong-Ying Li**, None; **Xi Hong,** None

## Notes

### Competing Interest Statement

The authors have declared no competing interest.

